# Lifespan associated global patterns of coherent neural communication

**DOI:** 10.1101/504589

**Authors:** Bikash Sahoo, Anagh Pathak, Gustavo Deco, Arpan Banerjee, Dipanjan Roy

## Abstract

Healthy ageing is accompanied by changes to spontaneous electromagnetic oscillations. At the macroscopic scale, previous studies have quantified the basic features, e.g., power and frequencies in rhythms of interest from the perspective of attention, perception, learning and memory. On the other hand, signatures and modes of neural communication have recently been argued to be identifiable from global measures applied on neuro-electromagnetic data such as global coherence that quantifies the degree of togetherness of distributed neural oscillations and metastability that parametrizes the transient dynamics of the network switching between successive stable states. Here, we demonstrate that global coherence and metastability can be informative measures to track healthy ageing dynamics over lifespan and together with the traditional spectral measures provides an attractive explanation of neuronal information processing. Finding normative patterns of brain rhythms in resting state MEG would naturally pave the way for tracking task relevant metrics that could crucially determine cognitive flexibility and performance. While previously reported observations of a reduction in peak alpha frequency and increased beta power in older adults are reflective of changes at individual sensors (during rest and task), global coherence and metastability truly pinpoint the underlying coordination dynamics over multiple brain areas across the entire lifespan. In addition to replication of the previous observations in a substantially larger lifespan cohort than what was previously reported, we also demonstrate, for the first time to the best of our knowledge, age related changes in coherence and metastability in signals over time scales of neuronal processing. Furthermore, we observed a marked frequency dependence in changes in global coordination dynamics, which, coupled with the long held view of specific frequency bands sub-serving different aspects of cognition, hints at differential functional processing roles for slower and faster brain dynamics.

## Introduction

A comprehensive understanding and characterization of the process of healthy aging are essential to treat age-associated neurological changes such as the decline in working memory, processing speeds, and executive cognitive functioning. Over the years, converging lines of evidence have successfully demonstrated the role of neural oscillations in many cognitive domains. Specific neuronal oscillatory patterns observed in EEG/ MEG data are essential markers of cognition (Buzsaki 2011), and researchers overwhelmingly agree on the use of field potential to tap neuro-cognitive processes associated with human brain function (Pesaran et al., 2018). Accordingly, several recent studies have tried to track age-related changes in the brain’s oscillatory profile using spectral estimation techniques. For example, the amplitude of resting and motor-related beta-band oscillations (16-25 Hz) is typically found to be higher in the older population compared to the younger population. Similarly, a substantial number of reports have highlighted that spontaneous peak alpha frequency (8-12 Hz) is lower in older people as compared to younger participants.

While age-related alterations in sensor specific features like power and frequency are fairly well-reported, very few studies have looked at changes in global patterns of frequency-specific synchronization in the context of healthy aging. We argue that much information about the mechanisms of aging is to be found in studying patterns of coherent activity across the lifespan. The relevance of this assumption can be assessed from the existing literature. For example, the theory of communication through coherence (CTC) posits that message passing across spatially distant neural assemblies demands coordinated fluctuations in their respective excitabilities (Fries, 2005). The importance of global coherence in the context of cognitive functioning is underscored by the essential need for efficient message passing in bringing about cognition. Even though the original CTC proposal was formulated in a task context, recent work has drawn out its repercussions for spontaneous brain dynamics, a.k.a resting-state activity (Deco et al.,2016). According to this formulation, resting brain activity frequently traverses across different functional configurations to maintain a state of maximal readiness in anticipation of external stimuli, which, when presented, collapses the state of the brain to whichever configuration is deemed most relevant in the stimulus context. In other words, resting-state brain activity must demonstrate metastable dynamics, whereby the brain fluidly recapitulates varied patterns of coherent activity (Deco et al.,2016). In line with this view, global metastability is found to be associated with cognitive flexibility and information processing in the brain. Therefore, tracking changes in coherence and metastability is crucial, given the fact that aging is marked by distinct cognitive changes that are, in turn, orchestrated by coherent neural oscillations.

The fundamental objective of this article is to track lifespan associated patterns of global coherence and metastability from neurophysiological recordings. As a necessary confirmation step, we first replicate the already well-established results in the field of aging neuroscience-the observations of reduced peak alpha frequency and an increase in average beta power with age on the current dataset. In doing so, we successfully establish the validity of earlier observations on a substantially larger dataset across the age continuum - a feature lacking in many previous studies. We then utilize a standard measure of global coherence to characterize band-specific lifespan trends. Finally, we apply a proxy for metastability - the standard deviation of the Kuramoto order parameter to characterize age-related alterations in frequency-specific metastable brain dynamics. Resting-state magnetoencephalogram (MEG) recordings from the Cambridge-Ageing Neuroscience (Cam-CAN) group for our purposes. Since this analysis has been carried out on a large cohort of an aging population (cross-sectional) consisting of 650 participants across an age range of 18-88 years we can consider them as a normative pattern of temporal structure of brain rhythms associated with ageing. The relevance of the global network metrics we further evaluated vis-à-vis performance in visual short term memory (VSTM) tasks over lifespan. Thus, we could summarize the organization of band specific coordinated brain dynamics over lifespan.

## Methods

### Participants

Cam-CAN is a multi-modal, cross-sectional adult life-span population-based study. The study was approved by the Cambridgeshire 2 Research Ethics Committee, and all participants have given written informed consent. The data presented here belonged to Stage 2 of the study. In Stage-1, 2681 participants had been home-interviewed and had gone through neuropsychological assessments and been tested for vision, balance, hearing and speeded response. Participants with poor vision (< 20/50 on Snellen test), poor hearing (threshold greater than 35 dB at 1000 Hz in both ears), past history of drug abuse, with any psychiatric illness such as bipolar disorder, schizophrenia, with neurological disease e.g. epilepsy, stroke, traumatic brain injury, or a score less than 25 in Mini-Mental State Examination were excluded from further behavioral and neuroimaging experiments. 700 participants had been screened from Stage 1 to Stage 2, of which Magnetoencephalogram (MEG) data from 650 subjects were available.

### Data acquisition

Data used in the preparation of this work were obtained from the CamCAN repository (available at http://www.mrc-cbu.cam.ac.uk/datasets/camcan/) (Taylor et al., 2016, Shafto et al., 2015). For all the subjects, MEG data were collected using a 306-sensor (102 magnetometers and 204 orthogonal planar magnetometers) VectorView MEG System by Elekta Neuromag, Helsinki, located at MRC-CBSU. Data were digitized at 1 kHz with a high pass filter of cutoff 0.03 Hz. Head position was monitored continuously using four Head Position Indicator coils. Horizontal and vertical electrooculogram were recorded using two pairs of bipolar electrodes. One pair of bipolar electrodes were used to record electrocardiogram for pulse-related artifact removal during offline analysis. The data presented here consisted only of resting state, where the subject sat still with their eyes closed for a minimum duration of 8 minutes and 40 seconds.

### Data preprocessing

Preprocessed data was provided by Cam-CAN research consortium, where for each run temporal signal space separation was applied to remove noise from the environment, from Head Position Indicator coils, line noise and for the detection and reconstruction of the signal from noisy sensors. All the data had been transformed into a common head-position. More details about data acquisition and preprocessing have been presented elsewhere (Taylor et al., 2017; Shafto et al., 2014).

### Data analysis

#### Welch spectrum

Fieldtrip toolbox (Oostenveld et al.,2011) was used to read the data provided in ‘.fif’ format. For each individual, data were downsampled from 1 kHz to 250 Hz. First, we sought to investigate age-specific changes in the spectral densities of the raw MEG signals.

Time series corresponding to the 102 magnetometers, resulted in a matrix *X* of size 102 × *T*, where *T* corresponds to the number of time points. Power spectral density for each sensor *c*’s time series *x*_*c*_(*t*)was estimated using Welch’s periodogram method. Each time series was divided into segments of 20 seconds without any overlap between segments. Spectrum was estimated for each segment after multiplying the time series segment with a Hanning window. Spectrums of all the segments were finally averaged.

We estimated a global spectrum, representative of each subject i.e. *S*_1_(*f*) by taking a grand average across the spectrums belonging to all magnetometers.

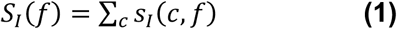

#### Quantification of spatial overlap between sources of alpha and beta activity in the sensor space

For each subject, the sensor map of alpha and beta activity were normalized separately.

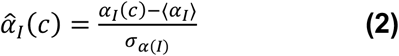

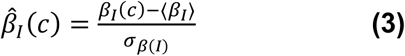

where *σ*_*α*(I)_ and *σ*_*β*(I)_ are the standard deviations of alpha and beta band respectively. Separation between the normalized sensor level representation *α*^^^_*I*_and *β*^^^_*I*_was indexed by the cosine angle between the two multidimensional vectors.

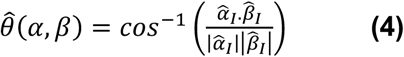

The angular separations across age were statistically analyzed using Spearman rank correlations and t-tests.

#### Global coherence

We calculated band-specific global coherence to measure the covariation of neural oscillations on a global level (Cimenser et al., 2008; Kumar et al.,2016). Global coherence among sensors at any frequency *f* is measured as the percentage of variance explained by the first eigenvector of the cross spectral density matrix at *f*.

In an individual subject’s data, for each sensor, the time series *x*(*t*)was divided into *N*non-overlapping windows of 5seconds duration each i.e. *y*(*t*). This resulted in an average of 112 (median, Interquartile range 1, range 70-220) windows for each subject. We employed 3 orthogonal discrete prolate spheroidal sequences (Slepian tapers) to avoid leakage in spectral estimates into nearby frequency bands. The time-bandwidth product was taken to be 2, which resulted in a bandwidth of 0.4Hz.

Before computing FFT, each data segment was detrended i.e. from each data segment *y*(*t*) the best straight line fit was regressed out.

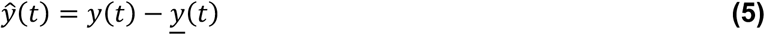

where *<u>y</u>*(*t*) is the straight line fit of *y*(*t*). Each segment was multiplied with a set of 3 orthogonal Slepian tapers and fast fourier transform was applied to the tapered segments.

Computing the complex FFT (for T tapers) at frequency *f* for each segment *n* of sensor *c* resulted in a complex matrix *Y* of dimension *F* × *C* × *N* × *T*. We utilized the chronux (Bokil et al., 2010) library to perform the global coherence analysis.

Cross spectral density between two sensors was estimated from Ŷ by using the formula

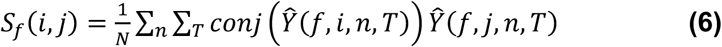

where *i*and *j* are the channel indices, *f* is the frequency index *n* is the segment index and T is the taper.

Singular value decomposition was applied to the cross spectral density matrix *S*_*f*_ for each frequency value *f*.

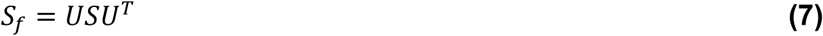

Diagonals of *S*would be proportional to the explained variance by the orthogonal set of eigenvectors *U*. The values of *S*were normalized so that each entry denote the percentage of the net variance explained in *S*_*f*_.

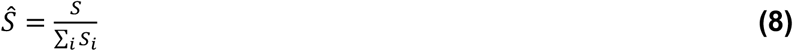

The first entry of *Ŝ* is defined as the global coherence. Global coherence was computed for each frequency value *f*, resulting an array *G* of length *F*.

#### Metastability

We calculated the metastability measure for all participants across all magnetometer sensors. Metastability is defined as variability of the Kuramoto Order parameter,*R*(*t*), which is given as,

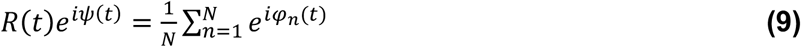

Where *φ*_*n*_ is the phase of the *n*^*th*^ oscillator and *ψ* is the mean phase of the system of oscillators. In this analysis, every MEG sensor is conceptualized as a coupled oscillator, summarized by its instantaneous phase *ϕ*(*t*). At any given point of time, the phase of each oscillator is extracted and projected onto a polar coordinate system, as a unit vector 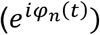. The length of the resultant vector, obtained from averaging all the unit vectors is interpreted as the Kuramoto Order parameter, *R*(*t*). The temporal variability of *R*(*t*) is measured by the standard deviation σ(*R*(*t*)), and defined as metastability (Deco et.al.,2017).

As a first step, the pre-processed resting state time series was band-pass filtered so as to obtain filtered time series. Instantaneous phase of each filtered band was estimated from the filtered data for metastability calculation. The pass band for the band-pass filtering step was kept narrow so that the resulting phase is readily interpretable.

As a first step, the pre-processed resting state time series was band-pass filtered so as to obtain filtered time series. Instantaneous phase of each filtered band was estimated from the filtered data for metastability calculation. The pass band for the band-pass filtering step was kept narrow so that the resulting phase is readily interpretable. For this analysis, each time series was filtered in the following frequency bands-2-4 Hz, 3-7 Hz, 8-12 Hz. Since valid phase estimation requires narrow pass bands, the beta band was further split into 2 sub-bands-16-20Hz and 20-25Hz and the respective metastability averaged. As mentioned earlier, the choice of frequency bands was dictated by phase considerations. An additional criterion was to chunk the frequency bands so that they map onto well-known frequency bands such as delta, theta, alpha and beta. As mentioned earlier, we restricted our analysis to below 40 Hz due to presence of HPI noise.

FieldTrip toolbox (ft_preproc_bandpassfilter.m) was used to band-pass filter each signal in the appropriate frequency bands. This routine was used to implement a finite impulse response (FIR), two-pass filter that preserves phase information of the time series.

Subsequently, instantaneous phase was estimated by using built-in MATLAB implementation of the Hilbert transform (hilbert.m). The resulting phase time series for each channel and participant was used to calculate band and subject specific metastability.

Similar to the preceding analysis, metastability analysis was performed by 1.) treating age as a continuous variable 2.) binning participants in the following age brackets - 18-35 years (Young Adults), 36-50 years (Middle Age), 51-65 years (Middle Elderly) and 66-88 years (Elderly).

For the region-wise analysis, the brain was segmented into 5 non-overlapping regions (frontal, centro-parietal, occipital, left and right temporal). Metastability index was calculated individually for all regions separately by randomly sampling 14 sensors from each region. Metastability was tracked as a function of age by calculating the Spearman rank correlations.

### Statistical Analysis

#### Continuous and categorical analysis of aging data

In order to bring all aspects of age-associated neural communication we performed both continuous and categorical analysis of the aforementioned brain measures with age as an explanatory variable. The primary goal of the continuous analysis was to capture the pattern change over lifespan (e.g., whether changes of the patterns are increasing/ decreasing). For this analysis we divided the whole cohort into bins of 5 years starting from 18 years. The bins were non-overlapping and the center of each bin was considered as the representative age value of the bin.

On the other hand, in the categorical analysis decomposing the whole data into cohorts with age ranges 18-35, 36-50, 51-64 & 66-88 allowed us to get finer and accurate insights in each stage of the adult span which has been well-documented in the fMRI literature (Chan et al., 2014) as well as the results obtained here can be contextualized with previous studies. The age ranges were unequally chosen because of the limitations posed by the CAM-CAN data set where different numbers of samples in each age group are available. However, in order to keep a reasonable number of samples > 120 in each cohort we chose the bins accordingly.

#### Regression analysis

Linear regression analysis was performed by separately considering each of the 5 estimated measures (Power, Global Coherence, Metastability, PAF and Topographical segregation Index) as response variables, while keeping age as the explanatory(continuous) variable.

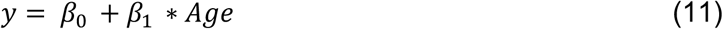

Linear regression was performed using the fitlm.m matlab procedure, which yielded an omnibus F-statistic, regression coefficients and goodness of fit(*R*^2^) and log-likelihood(L). The regression coefficient was taken to represent effect size. Additionally, we also considered 2nd and 3rd order polynomial fits such as-

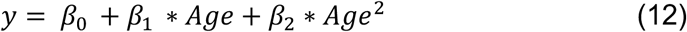

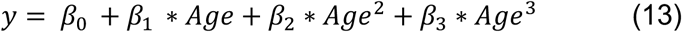

Akaike Information Criteria was used for model selection and was calculated as-

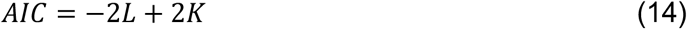

Where K is the number of model parameters including the intercept.

For the analysis that report spearman correlations, Effect sizes were computed using cohen’s d

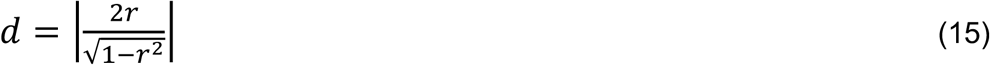

(Reported in Supplementary Material in detail)

#### Categorical Analysis

To systematically evaluate the relationship between age and brain measures Power, Coherence, metastability, PAF and Topographical segregation we used Spearman correlation analysis following what was described at Khan et al., 2018. Except for the logarithm of power, the other measures are not guaranteed to be following a Gaussian or normal distribution, hence a common non-parametric test, Spearman correlation was chosen to evaluate all correlations in this article.

In brief, pairwise comparisons between groups were performed using permutation testing. In each iteration the groups were collapsed and random draws were made to form random groups. Difference of means was calculated for the random group assignments and the procedure repeated for 10000 iterations to construct a surrogate distribution. Finally, significance was estimated using the surrogate distribution. The statistics were reported in terms of p-values, effect sizes and difference of means.

## Results

Our analysis strategy was two-fold. First, we conducted a categorical analysis by chunking the age-continuum into discrete groups (see Table 1). We have divided the age values of total N=650 subjects into four age groups: Young Adults (YA), Middle Elderly (ME), Middle Late (ML), Older Adults (OA), for which the demographic information has been provided in **Table 1**. Earlier studies have done similar grouping (Chan et al 2014) and care was taken such that we have sufficient number of participants in each age group > 120. Subsequently, we considered age to be a continuous variable with bins consisting of 5 years between 18-88 and performed linear and polynomial regression to estimate age associated trends. The bins were non-overlapping and the center of each bin was considered as the representative age value of the bin.

**Table 1.** Sample size and gender statistics in each representative age group

| Age group | N | % Female |
| --- | --- | --- |
| YA (18-35) | 126 | 55 |
| ME (36-50) | 159 | 49 |
| ML (51-65) | 149 | 50 |
| OA (66-88) | 216 | 46 |

### Age trajectories in MEG resting state brain dynamics

We studied the effect of healthy ageing on the fundamental properties of the endogenous band-limited neural oscillations such as amplitude and center frequency. Since the Head Position Indicator (HPI) coil related noise can be unreliable at higher frequencies, we concentrated our analysis between 0-40Hz which fully contains the neural oscillations in the **Delta** (1-3Hz), **Theta** (4-8Hz), **Alpha** (8-12Hz), and **Beta** (16-25Hz) frequency bands.

#### Band limited power

An omnibus ANOVA that considers age as the explanatory variable yielded significance for all measures tested (**details reported in Supplementary material**). Our analysis revealed that spectral power in **Delta** (*β* = 0.008, *p* = 0.1), **Theta** (*β* = 0.004, *p* = 0.46) and **Alpha** (*β* = 0.0002, *p* = 0.76) bands did not vary with age (**see Fig 1**). In contrast, beta band power varied significantly with age (*β* = 0.03, *p* < 0.0001). Visual inspection of **Fig. 1d** seemed to suggest 18-22 to be an outlier group, so we additionally removed that group and compared linear and 2nd order polynomial fits for the beta band and found the 2nd order polynomial fit to be better (AIC linear = 2722, AIC quadratic = 2718). Alpha band power was estimated by averaging the estimated spectral values within 8-12 Hz. We further quantified age-effects in the beta band using a categorical approach and found significant differences in group means between YA vs ME, YA vs ML, YA vs OA and ME vs OA groups.

**Figure 1.**
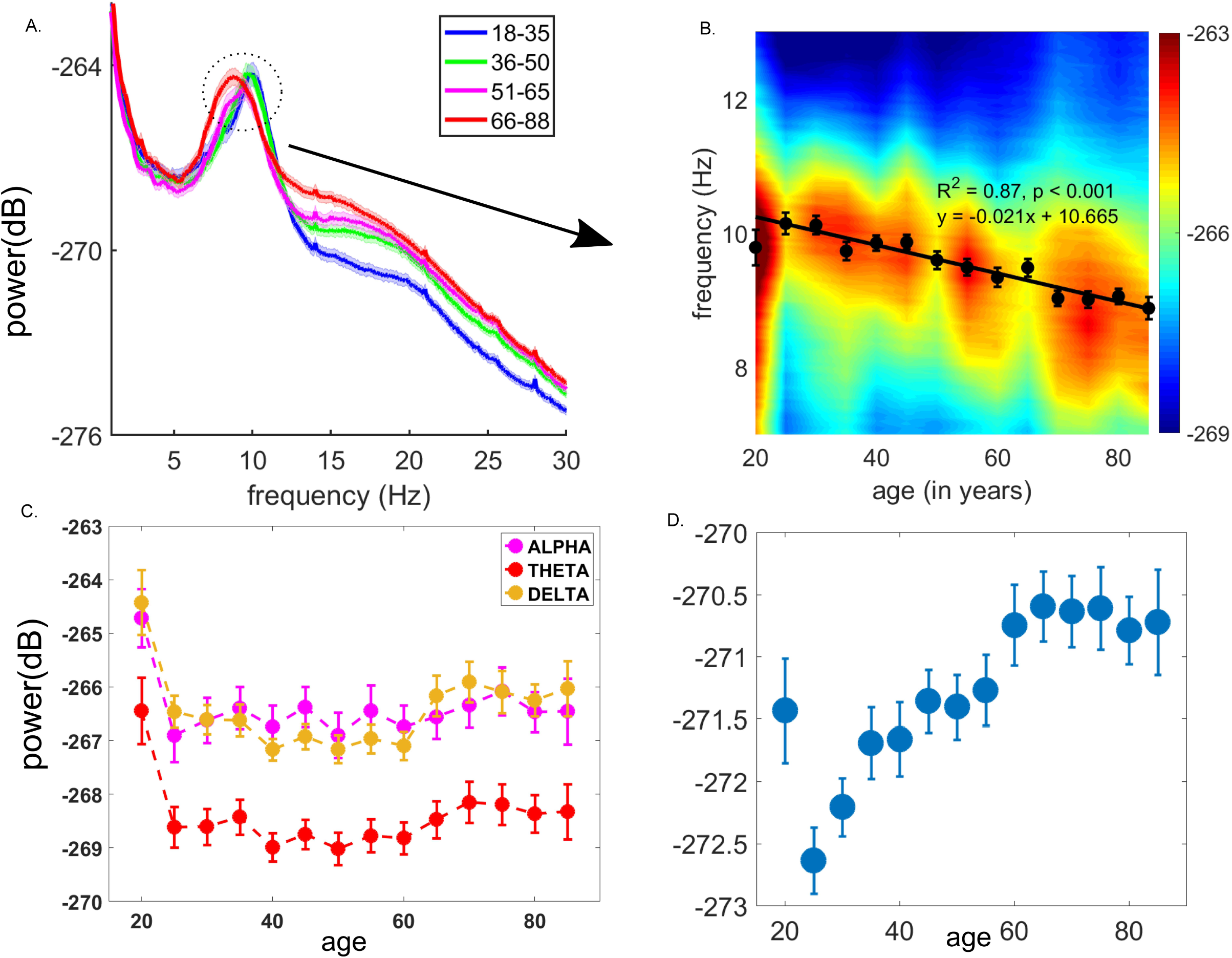
Relation between global spectral activity and age. **A**. Plots of mean power spectral density for 4 non-overlapping age groups i.e. 18-35, 36-50, 51-65 and 66-88. Shaded region denote standard error of mean. **B**. Variation of alpha activity with aging. Center frequency in the alpha band for each age bin has been plotted as solid circles and solid black line is the linear fit of these points (labels indicate effect sizes, significance and correlation function) **C**. Spectra in the delta, theta and alpha bands as a function of age. **D**. Beta spectra as a function of age

#### Peak alpha frequency (PAF) shifts with age

The center frequency in the alpha band (8-12 Hz) has long been studied in the context of healthy and pathological aging (**see Fig 2**). Here, we sought to quantify age related variations in PAF by averaging the center frequency across the 102 sensors and correlating the global mean with age as a variable. The regression analysis confirmed age-related reduction in PAF (*β* = −0.01, *p* < 0.0001). Frequency value at which there was maximum activity in the alpha band i.e. 8-12 Hz for a subject, was taken to be the peak alpha frequency of that subject. **Fig 1B** is an age-spectrogram which shows variation in the power spectral density in the alpha band with age.

**Figure 2.**
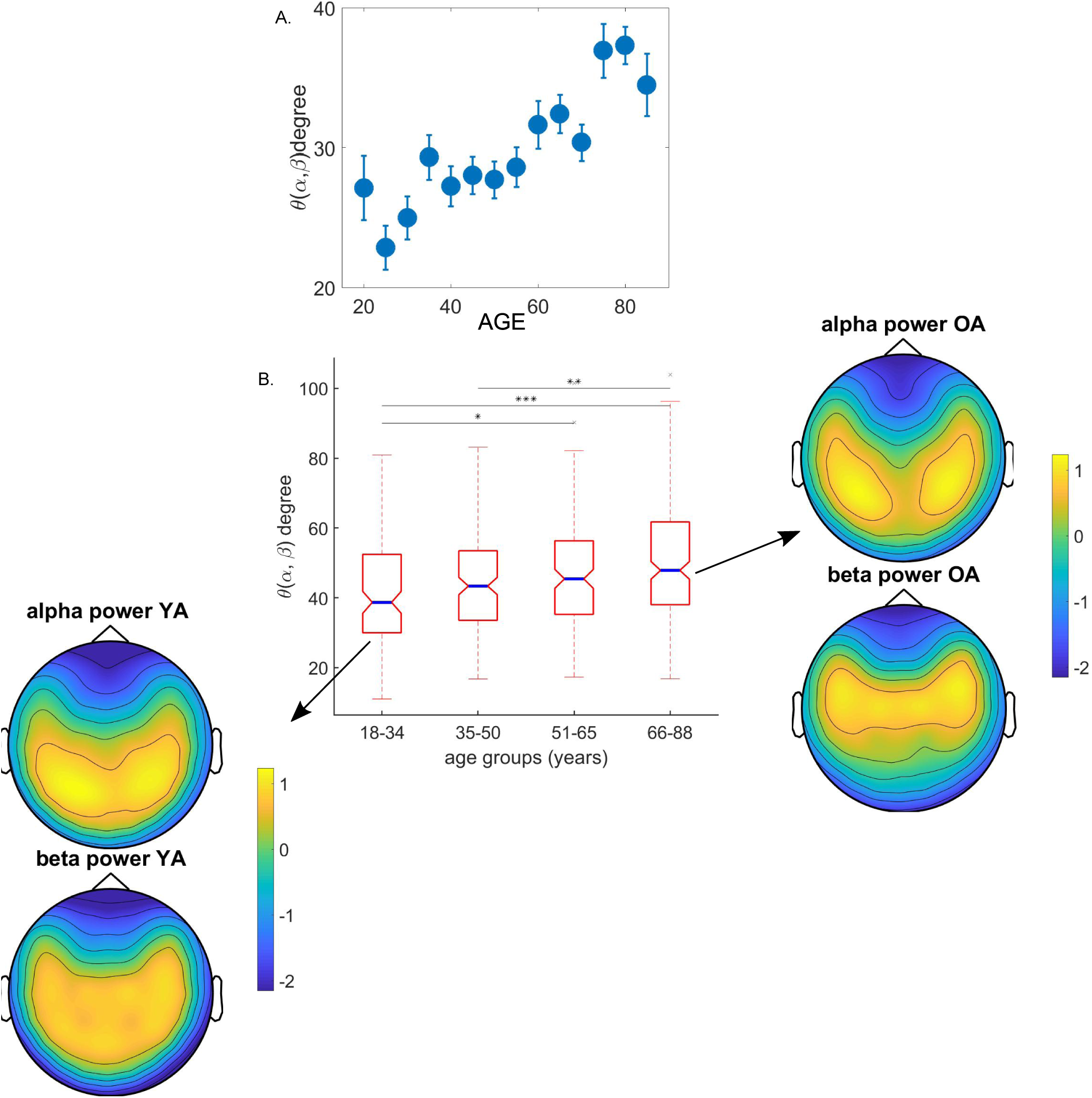
Segregation of sensor level topographies with aging. **A**. Angular separation between alpha and beta bands(in radians) as a function of age. **B**. Boxplot for the distribution of angles between the sensor topographies of center alpha power and average beta power for the four age groups. Blue line denotes the median of the distribution and the notch indicates 95% confidence interval of the median. Inset:Sensor topographies of alpha power at center frequency and average beta power for the two extreme age groups.

Next, we split the age-range into discrete categories and performed permutation tests to estimate group differences across age groups. YA was found to differ significantly with ME, ML and OA whereas the ME group differed from OA in terms of the sample means (effect sizes reported in Supplementary Materials).

#### Topographical distribution alpha and beta band power

Next, we investigated the spatial topographies of whole brain networks corresponding to age-related difference along slow and fast time scales of neuronal signal using subspace analysis borrowed from linear algebra. We quantified the overlap between the two sensor topographies by the angle between their respective vector representations (**See Fig 2**). Larger angles indicated more separation and less topological overlap between sensor groups. The topographical separation between the sensor−wide distribution of alpha and beta power was found to increase with age (β=0.003, *p* < 0.0001) (**Fig 2A**). **Fig 2B** shows the average topographical map of alpha activity at the center alpha frequency and average beta activity in 16-25 Hz for the youngest and oldest age groups.

Although we observed similar patterns of difference between the oldest and youngest age groups for global alpha band power and beta band power, there seemed to be a qualitative difference in the overlap of sensors representing alpha and beta band activity. The categorical analysis revealed that sample means in YA group differed from ME, ML and OA. ME differed from ML and OA whereas ML was different from OA (effect sizes reported in Supplementary Material).

### Age trajectories in band-specific global network measures: coherence and metastability

#### Global Coherence over lifespan

Presence of large-scale functional brain networks was investigated using global coherence across all MEG sensors at different frequencies for each subject (Cimenser et al,2011; Kumar et al. 2016). Whole-brain coherence was summarized as the ratio of the principal eigenvalue to the sum of all eigenvalues of the inter-sensor coherence matrix (see Materials and methods).

Global coherence at all the frequency values within a frequency band were averaged to generate a representative value for the corresponding frequency band in four age groups, YA, ME, ML, OA (**Fig 3A)**. Representative global coherence in age bin was averaged for the continuous analysis and standard error was computed for each age bin (**Fig 3B**). Global Coherence in the delta and theta band was found to increase with age – delta (β=0.0005, *p* < 0.0001), theta (β=0.0002, *p* = 0.009). In contrast, global coherence in the alpha band varied inversely with age (β=-0.0008 *p* < 0.001) while Beta band global coherence did not display an age effect (β∼0, *p* = 0.86) (See **Fig 3B)**.

**Figure 3.**
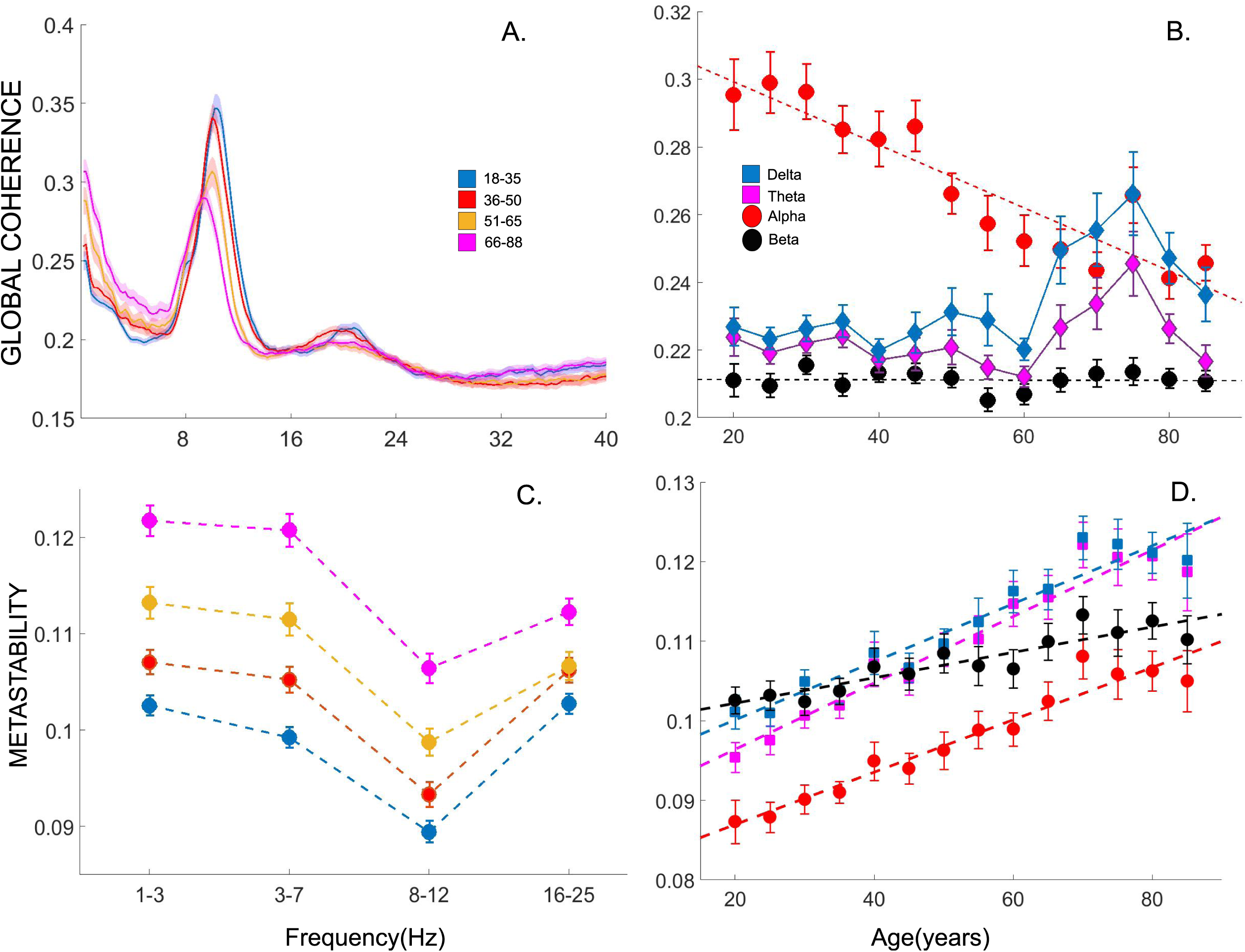
Differential changes in global coherence with aging. **A**. Plots of mean global coherency for the four age groups. Shaded region denotes s.e.m. **B**. Differential variation of global coherence for frequency bands. **C**. Metastability for four age groups in delta, theta, alpha and beta bands.

#### Metastability and aging

We estimated the variability of neuronal communication states using metastability as a function of age and frequency. We observed a dichotomous pattern in metastability as a function of frequencies in all age groups - a sharp decrease with increasing frequencies till 12 Hz and a gradual increase in the metastability indices across frequencies between 12 – 40 Hz (**Fig 3C**). Qualitatively, we found metastability to be higher for delta, theta and beta bands as compared to the alpha band. Interestingly, in all age categories, the variation of metastability with frequencies was consistent, essentially a U-shaped profile. From the continuous analysis we could establish that band-specific metastability increased with age across all frequency bands-Delta (β=0.0004 *p* < 0.0001), Theta (β=0.0004 *p* < 0.0001), Alpha (β=0.0003, *p* < 0.0001), Beta (β=0.0001, *p* < 0.0001).

#### Region-wise analysis of metastability reveals differential trends

In order to track changes in metastability in specific brain areas we segmented the sensors in 5 groups - Frontal, Centro-Parietal, Occipital, Left Temporal and Right Temporal regions. The region-wise analysis consisted of 14 randomly sampled sensors to compute metastability in each brain region. Next, we tracked the region-wise metastability with aging. Spearman rank correlation was performed to characterize trends in band and region specific metastability and effect sizes quantified using Cohen’s d. Delta and Theta oscillations either stayed invariant or reduced as a function of age in the occipital, left temporal and right temporal regions. Beta band metastability showed the highest age correlation (using Spearman rank test) in the centro-parietal sensors while staying invariant in the occipital and temporal sensors. The following Spearman rank coefficients and effect sizes (Cohen’s d) were obtained for delta band - frontal (*c* = 0.2499, *d* = 0.249, *p* < 0.001), centro-parietal (*c* = 0.1629, *d* = 0.3302, *p* < 0.001), occipital (*c* = −0.0686, *d* = −0.13, *p* = 0.0805), left temporal (*c*∼0, *p* = 0.92), right temporal (*c* = −0.1428 *d* = −0.28 *p* < 0.001). The corresponding values for alpha band were frontal(*c* = 0.2161, *d* = 0.44 *p* < 0.001), centro-parietal (*c* = 0.1808 *d* = 0.36, *p* < 0.001), occipital (*c* = 0.1348 *d* = 0.27, *p* < 0.001), left temporal (*c* = 0.2030, *d* = 0.414 *p* < 0.001), right temporal (*c* = 0.2070, *d* = 0.42, *p* < 0.001). For theta band-frontal (*c* = 0.1725, *d* = 0.35, *p* < 0.001), centro-parietal (*c* = 0.2049, *d* = 0.416, *p* < 0.001), occipital (*c* = 0.1141, *d* = −0.2, *p* < 0.001), left-temporal (*c* = −0.04 *d* = −0.08 *p* < 0.2396), right-temporal (*c* = 0.0457, *d* = −0.08 *p* < 0.2443) were obtained. We tracked metastability in the beta band for two frequency bands (β_1_, β_2_) using similar statistical methodology. Centro-parietal sensors showed the highest age-related positive correlations (*c* = 0.2046, 0.2542, *d* = 0.418 0.51, *p* < 0.001 for the two bands respectively) (Fig 4).

**Figure 4:**
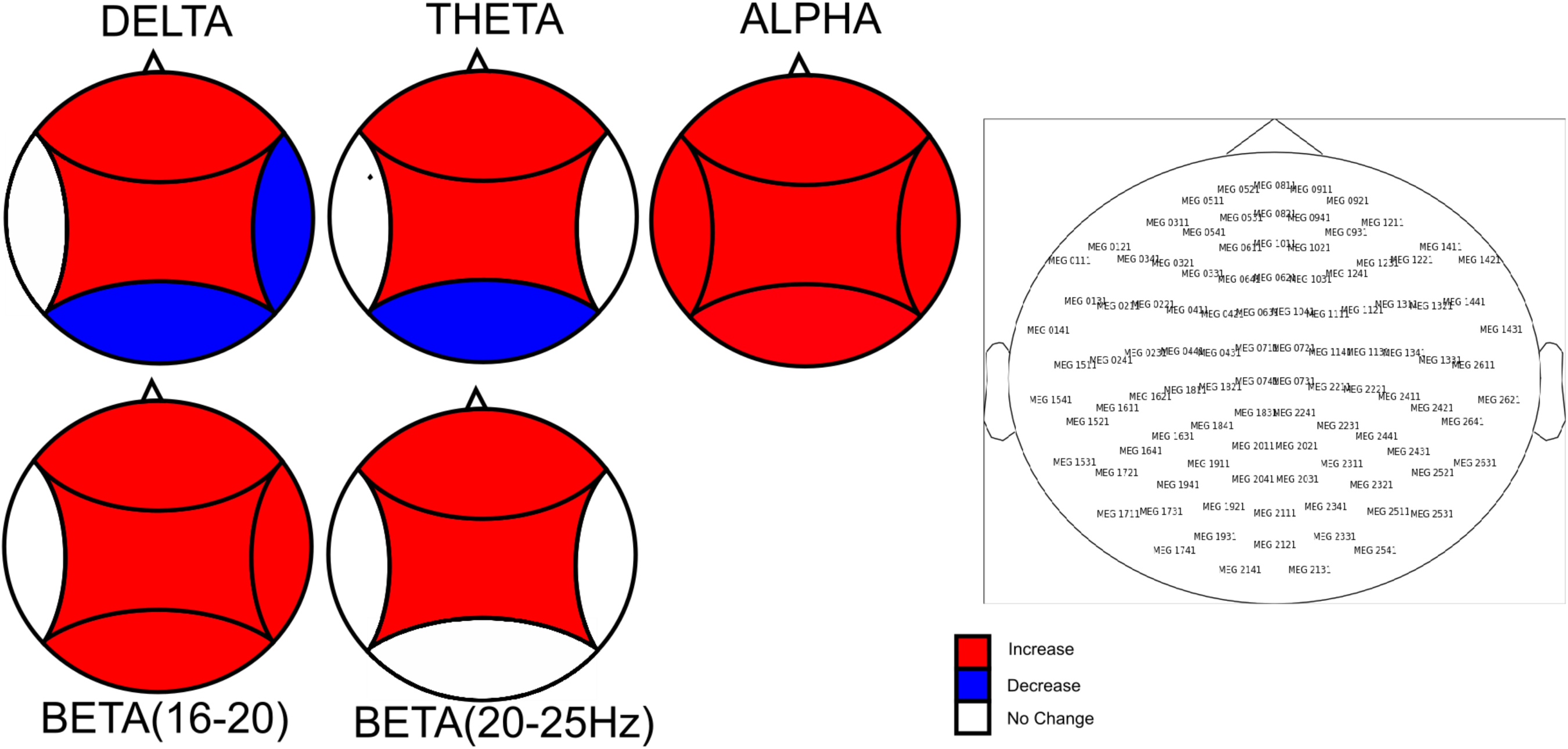
Region-wise increase and decrease in global metastability. **A**. Shows the results for the region-wise metastability analysis. Colors indicate the direction of the age-related trend as measured by the spearman rank correlation coefficients. 14 sensors were chosen at random from each of the 5 anatomical areas-frontal, centro-parietal, occipital, left and right temporal. **B**. Vector View magnetometer layout.

#### Relationship of between global network measures and performance metrics over lifespan

In order to evaluate the relationship of normative brain rhythms over lifespan we computed the correlations between global network measures and the performance metric of precision in a visual short-term working memory (VSTM, previous used by Zhang et al 2008) task available with the Cam-CAN cohort. We observed a significant correlation at 95% confidence levels between global coherence in the alpha band and precision in VSTM task after regressing out the effect of age (*ρ* = 0.09, *p* = 0.0143, **Fig 5**). The global coherences and metastability computed in other frequency band were not significantly correlated with precision (**detailed statistics reported in Supplementary Materials**).

**Figure 5:**
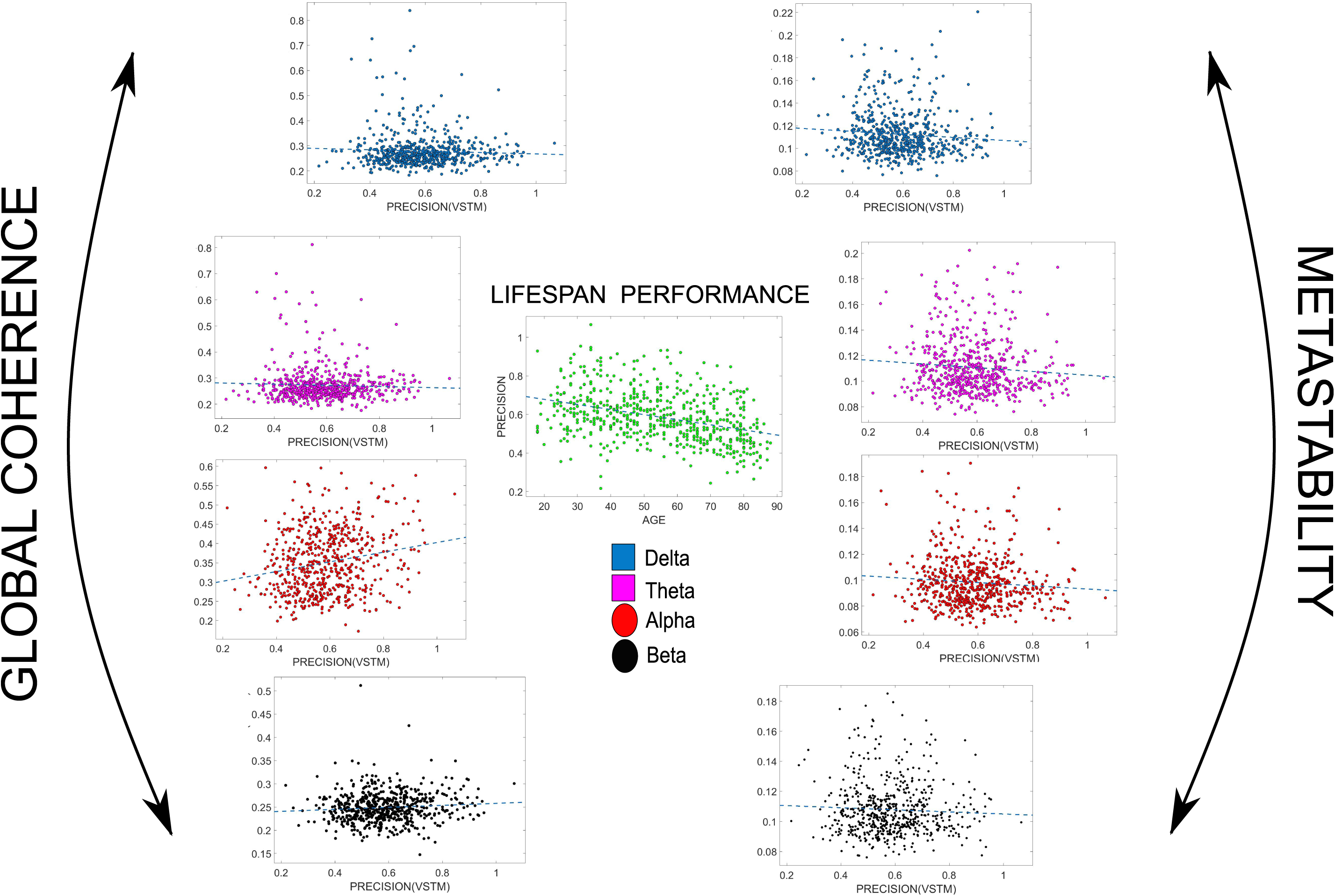
Correlation of VSTM precision with global coherence and metastability. **Center:** Scatterplot of precision with age. **Left:** Scatterplot of band-specific global coherence with precision in VSTM task. **Right:** Scatterplot of band-specific global coherence with precision in VSTM task.

## Discussion

Neuronal communication is the backbone of basic human brain functions and supports a myriad of cognitive functions at various scales of nervous system organization (Mesulam 1990). While spectral estimates attempt to link neural oscillations with cognition (Henry et al. 2016, see Pesaran et al. 2018 for a nice review), very few studies are available that provide normative mapping of neuronal oscillations across healthy lifespan aging. We observed a significant age-related decline in peak alpha frequency (PAF) at the sensor level as well as increase in broadband beta power in a healthy cohort consisting of 650 human participants from the CAMCAN repository (Fig 1). Out of all sensor specific spectral features such as frequency and amplitude of oscillations in narrowband and broadband, PAF and beta power varied exclusively with age in opposite ways (Fig 1). Subsequently we could track the global subspace that sculpts the alpha and beta topographies, and their corresponding overlap over lifespan. Interestingly the angular separation between the alpha and beta topographies increase with age indicative of segregation of underlying generators over lifespan (Fig 2). Integrative mechanisms operational at the macroscale of whole-brain MEG sensor-space were captured by two complementary mathematical frameworks – global coherence spectrum that parametrizes the strength of band specific synchronization in different frequencies over a set of network nodes (MEG sensors for the purpose of this paper), and metastability that captures the degree of intermittency that exists between two successive synchronization states (Fig 3). Together, the two measures along with the spectral estimates quantifies the dynamic repertoire of the state variables. We observed an emerging dichotomy with aging in pattern of global coherence across slow and fast time scales (Fig 3A, B). The global alpha coherence decreases over lifespan followed a linear relationship whereas global beta coherences are unaffected by aging. The global coherences in slower frequencies - delta and theta on the other hand, are unaffected by aging up to a critical age of around ∼45-50 years (Fig 3B). Thereafter, the global coherence shows an increase with age up to ∼70 years and decrease further upon reaching a peak value. While alpha global coherence could be fitted best with a linear curve, theta and delta variation over age was non-linear. Concurrently, metastability exhibited a monotonically increasing relationship over lifespan in all frequencies (best fitted by a linear curve), with a visible saturation for elderly (∼70 years) (Fig 3C, D). Interestingly, while increase in metastability in alpha band was truly global, increases in metastability in other bands were region-specific (Fig 4). Furthermore, while global coherence may be reflective of task performance, metastability, which is essentially a measure of phase variability was uncorrelated with task performance (Fig 5). In summary, we present a comprehensive account of the temporal properties of neuro-electromagnetic signals over lifespan aging that can be further interpreted in relation to recent and established findings in the literature to develop a neurodynamic explanation of several important observations in healthy aging (Fig 1-5). Furthermore, we argue that our results are extremely helpful for understanding the pathological aging scenarios (e.g. Alzheimer’s and Dementia) beyond the standard model of “accelerated ageing” (Toepper 2017)

The decrease in PAF, prominently observed in our study, has been reported to be a biomarker of normal and pathological aging process, especially for dementia, mild cognitive impairment, and Alzheimer’s disease (Scally et al., 2018; Dickinson et al., 2018; Osipova et al., 2005; Jeong, 2004). Patients with Alzheimer’s disease show a significant decrease in PAF compared to age-matched control group (Osipova et al., 2005; Jeong, 2004). Parkinson’s patients with dementia have a lower PAF compared to age-matched controls (Soikkeli et al., 1991). Interestingly, developmental changes in spontaneous electrocortical activity is associated with an increase in PAF from early to late childhood (Miskovic et al., 2015). While the mechanistic explanation of this dichotomy still remains elusive, several computational attempts have suggested a link between the thalamocortical circuitry responsible for alpha rhythmogenesis and age-related morphological differences in thalamocortical circuits to explain slowing down of PAF. In fact, PAF may carry the signature of an ending of rapid neurodevelopmental process of human beings, behaviorally observed as trait developments from adolescence to young adults. Concurrently, cognitive task relevant EEG/MEG studies have linked PAF with scores on cognitive paradigms such as working memory (Clark et. al 2004) and visual acuity (Samaha et. al 2015) suggesting a crucial role of PAF with age associated changes in attention and memory from YA to OA. Consistent with extant literature, power in the beta band was found to increase with age. Increase in the band-limited beta power in older population compared to younger population has been reported both in the context of resting state and sensory-motor tasks (Rossiter et al., 2014; Heinrichs-Graham et. al, 2016), where beta oscillations have been regarded as an index of motor inhibition and volitional movement (Heinrichs-Graham et. al 2016). Subsequently, we depart from some earlier studies in key respects. Firstly, we find that band-limited power (spectral feature independent of frequency in our study) in the delta, theta and alpha bands does not vary with age. Second, the angular overlap decrease between the alpha and beta bands topographies with age reflects a segregation of function which can emerge from neuro compensatory functional mechanisms or possibly structural decline e.g., myelination degradation. Neurodegenerative pathologies like AD and Parkinson’s share many similarities with healthy aging, due to which many have speculated whether neurodegeneration is an accelerated aging process (Toepper 2017). Remarkably, the two sets of features, PAF and band-limited beta power exhibits different age associated trajectories. While decline of PAF is best described with a linear model, the band limited beta power trend is best described by a quadratic curve across age continuum. Thus, our results throw in the possibility that while PAF decrease observed in pathological scenarios may be a non-specific marker of disease, other features like beta power increase may be more relevant candidates to tag preservation of function via neuro compensatory mechanisms.

A key contribution of our study is the archival of global network measures over lifespan, particularly that are relevant for the neural information processing time scales. Few recent studies using M/EEG have further emphasized that patterns of age-dependent segregation for beta and gamma mediated networks differed substantially during maturation (Miskovic et al., 2015; Khan et al., 2018). A recent study by Khan et al., 2018 further reports that beta band mediated networks become more locally efficient, i.e. tending towards clustering and more connections with adjacent regions with age, while gamma band mediated networks become more globally efficient, i.e. tending towards shorter overall path lengths and thus faster communication across larger cortical distances, with age during maturation. However, how do such large-scale and local communication organize and orchestrate across different sensors and in different bands during various stages of healthy adult lifespan remains largely unknown. In our study, we attempt to quantify the band specific normative values as features during resting state borrowing the concept from Communication Through Coherence (CTC) hypothesis. CTC operationally defines neuronal communication as generation of coherent activity across neuronal assemblies. This view holds that interareal coherence presents windows of excitability where communication channels between brain regions are maximally utilized (Fries, 2005). Resting state brain activity is said to reflect the brain’s tendency to engage and disengage these channels of communication spontaneously (Deco et. al, 2011). From a dynamical systems perspective, spontaneous brain activity must exhibit metastable brain dynamics, whereby the global brain dynamic stays clear of the two extremes of constant synchronization and desynchronization and instead, periodically shuttles back and forth between coherent and incoherent regimes. More formally, global coherence indexes the average phase and amplitude correlation across sensors whereas metastability measures the variability in phase relationships of sensors across time. The complimentary, yet related nature of global coherence and metastability offers unique insights into the mechanistic underpinnings of global brain dynamics. An example of this is a recent computational study by Vasa et al. which describes how local lesioning in nodes with high eigenvector centrality leads to a simultaneous decrease in global synchrony along with an increase in metastability (Vasa et al., 2015). For a review of the complementary nature of global coherence and metastability, see (Hellyer et al. 2015, Vasa et al. 2015, Deco et al. 2017). The global coherence decreases in alpha band (8-12 Hz) with concomitant increase of metastability over lifespan indicates the transformative role of alpha over the aging process. This would strongly suggest that alpha can be a possible mode of neural communication associated with neural compensation whereas slower frequencies like delta and theta may reflect a critical juncture in adult lifespan at certain age ranges. Interestingly the critical age ranges from where delta and theta global coherence start peaking (∼45-50) is a critical phase of life in terms of performance where ability to learn new skill starts diminishing (Janacsek et al. 2012). We argue while alpha coherence decrease may be associated with neuro compensatory mechanisms, they may not have a direct bearing on performance for which delta and theta may be more informative. Studies have demonstrated that theta rhythms are crucial for information processing underlying sequence learning (Sauseng et al. 2009, Koene and Hasselmo 2009) which is clearly a relevant metric for skill learning observed by Janaccsek et al. 2012.

Our earlier work has proposed that a way to implement the CTC hypothesis, that is, optimal exploration of the dynamical repertoire inherent in the brain structural connectivity, is by maximization of metastability (Deco et. al., 2016). Here, we interpret metastability as a measure of the variability of the states of phase configurations with time. Thus, metastability should decrease with the introduction of external stimulation and task conditions. In terms of dynamical systems, resting brain to exhibit maximal metastability, refining and providing evidence in favor of the synergetic hypothesis of Haken (Corning 1995) (later further explored by Tognoli and Kelso 2014). We observe an age-related increase in global metastability across all frequency bands and the trend is best fitted by a linear model. This result can be contextualized from two opposing theories of healthy aging. The method of neuro-compensation argues that age-related changes in brain dynamics suggest a compensatory mechanism by which function gets restored in response to structural decline (Naik et al. 2017). In this regard, it is interesting to note that Alzheimer’s disease and traumatic brain injuries (TBI) are associated with a reduction in global metastability (Córdova-Palomera et al., 2017; Hellyer et al., 2015). Since metastability is a direct measure of the functional capacity of the brain and has been shown to confer cognitive flexibility in task-switching, information-processing and logical memory (Hellyer et al. 2015), this would argue in favor of a compensatory explanation of the global increase in metastability with aging. However, the neural noise hypothesis of aging would suggest a different interpretation. This theory argues that age-related cognitive decline is best explained as a consequence of an increase in the noisy baseline activity of the brain (Voytek et al., 2015; Dave et al., 2018). According to this framework, global phase inconsistencies as we observe here is an obligatory change resulting from change in underlying scaffold dictated by gradual change in white and grey matter volume that shifts the baseline and result in an unspecific lifespan-associated increase in neural noise. Within this framework, changes in global metastability and coherence reflect an epiphenomenon that occurs due to an increase in neural noise. Future efforts should focus on resolving this debate. One possible direction would be to study brain signals through measures of signal complexity using source reconstructed EEG/MEG, to elucidate the role of specific brain regions in bringing about metastable patterns of activity. More direct estimates of metastable state switching from electrophysiological data could be employed to disentangle the effects of noise. Recent works in this direction have proposed ways to directly estimate metastable switching between synchrony states. For example, Vidaurre et.al. 2016 propose a Hidden Markov Model (HMM) based method to decompose electrophysiological time series into recurrent, quasi-stationary phase-locked regimes. This involves fitting source reconstructed time series with multivariate autoregressive models and modelling state switches through the HMM approach. Another promising avenue would be to invoke whole brain computational models which incorporate neural plasticity mechanisms that operate at time scales that are relevant to aging (Vattikonda et al., 2016; Abeysuriya et al., 2018). This is also necessary to reconcile the region specific metastability patterns we observe across frequency bands, with alpha band metastability increase being truly global versus region-specific enhancement and decrease of metastability in other frequency bands over lifespan (Fig 4).

An ongoing research direction in the neuroimaging community is to relate resting state dynamics to performance measures also sometimes referred to as behavioral phenotypes (Nomi et al 2017, Liegeois et. al 2019). While the slow time-scale of fMRI has been primarily used for this purpose to argue about cognitive flexibility from resting state functional connectivity (FC) metrics (Naik et al 2017, Nomi et al 2017, Liegos et al 2019), the variation of the global coherence and metastability from MEG presented us an opportunity to investigate the relationships between global network properties and task performance in the neural times-scale. The CAM-CAN dataset has the Verbal short-term memory task (VSTM) in which the accuracy is anti-correlated linearly with increase in age. Interestingly only the global coherence in alpha band was correlated with precision when the age effects were corrected, while the global coherence in other frequency bands are uncorrelated with VSTM performance (Fig 5). On the other hand, metastability has no bearing on performance accuracy once the effects of ageing were considered for any frequency. Thus, except the global network captured by alpha global coherence, the others are non-specific measures of neurophysiological processing. In other words, measures like global coherence/ metastability quantifies the overall shift global information processing rather than being relevant for a specific task.

An important caveat of our study was due to limitation posed by the Cam-CAN data set particularly because of the presence of harmonics of lower frequencies being present in higher frequencies, a systematic analysis of gamma band was not possible. The gamma frequencies in the resting state did not show any statistically significant differential change with ageing at least in the low gamma range (30-40 Hz), although the task data showed interesting patterns. However, such discussions remain out of scope of this paper. Another limitation is that in spite of large sample size the current analysis is restricted to the sensor level, our results are so far only indicative of activity at the neural level. Source reconstruction may provide a direct estimate of global coherence at the level of neural assemblies and help in elucidating its relationship with sensor level global coherence and metastability, the efforts toward which will be presented elsewhere in future. Caution is also required in interpreting the results due to the modest effect sizes involved in certain measures. Similarly, caution is warranted in interpreting global coherence and metastability measures. Due to the way it is constructed, global coherence may give misleading information under some circumstances. For example, it is possible to obtain spuriously high values of global coherence even when the underlying signals are independent when most of the power is concentrated in a few sensors. In this study, the almost evenly distributed scalp topographies (Fig.3) would preclude that possibility. Future work is underway to address some of these issues and limitations. The other limitation of the present study is the use of simple statistical models to explain spectral features as a function of age, but more complicated component models can be used in future (e.g., in Liegeois et al. 2019). A multidimensional analysis by estimating FC dynamics corresponding to different performance measures using big data techniques can further shape the understanding of rest and task

## Acknowledgements

This study was supported by NBRC Core funds, Ramalingaswami Fellowships (Department of Biotechnology, Government of India) to DR (BT/RLF/Re-entry/07/2014) and AB (BT/RLF/Re-entry/31/2011) and Innovative Young Biotechnologist Award (IYBA) to AB (BT/07/IYBA/2013). AB also acknowledges the support of the Centre of Excellence in Epilepsy and MEG. DR was also supported by SR/CSRI/21/2016 extramural grant from the Department of Science and Technology (DST) Ministry of Science and Technology, Government of India. Data collection and sharing for this project was provided by the Cambridge Centre for Ageing and Neuroscience (CamCAN). CamCAN funding was provided by the UK Biotechnology and Biological Sciences Research Council (grant number BB/H008217/1), together with support from the UK Medical Research Council and University of Cambridge, UK. In accordance with the data usage agreement for CAMCAN dataset, the article has been submitted as open-access.

---

**Fig S1.**
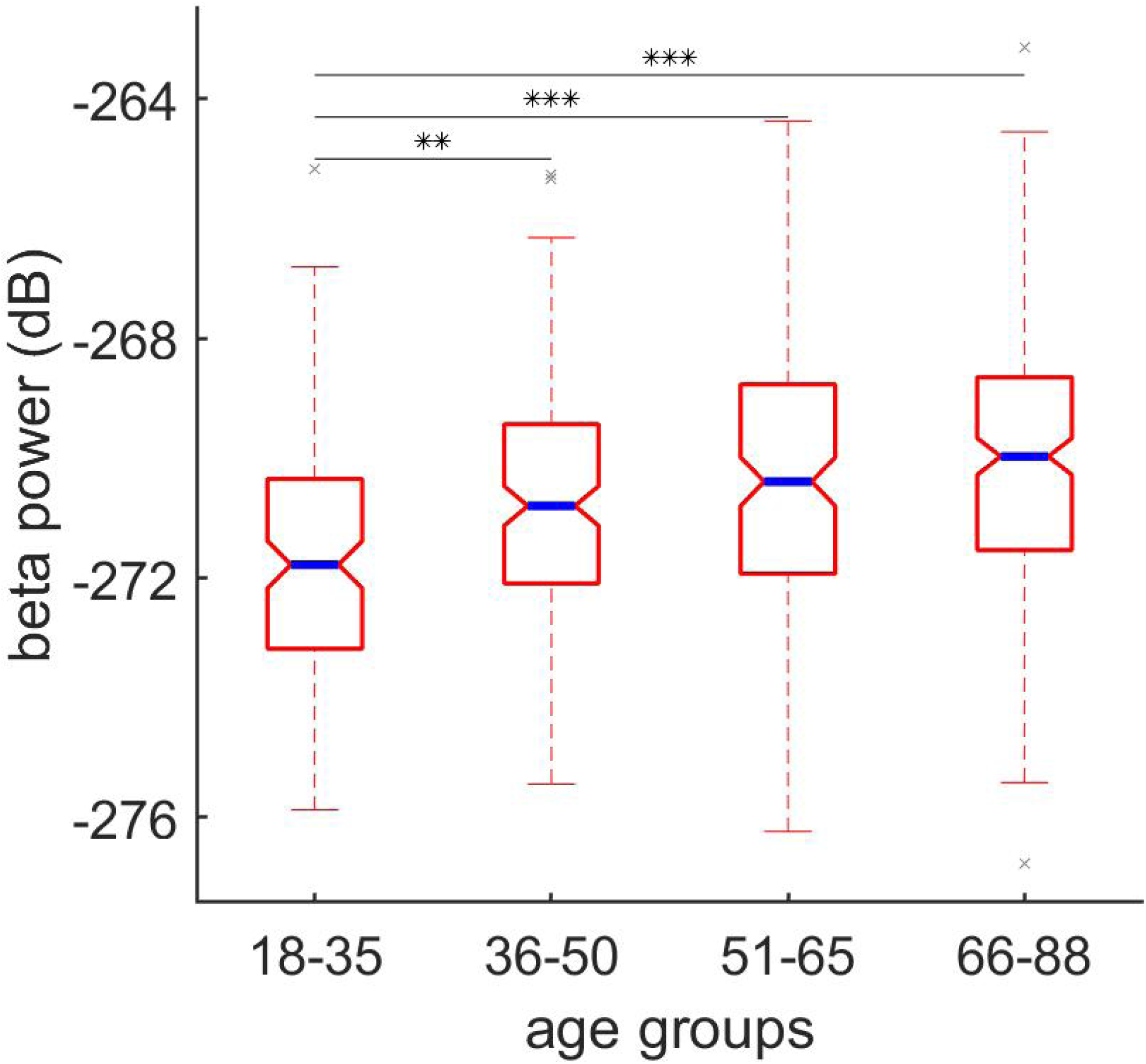
Boxplot of distribution of band limited power in the beta band (16-25 Hz). Blue line indicates the median of each distribution. Notch denotes 95% confidence interval of the median

**Fig. S2.**
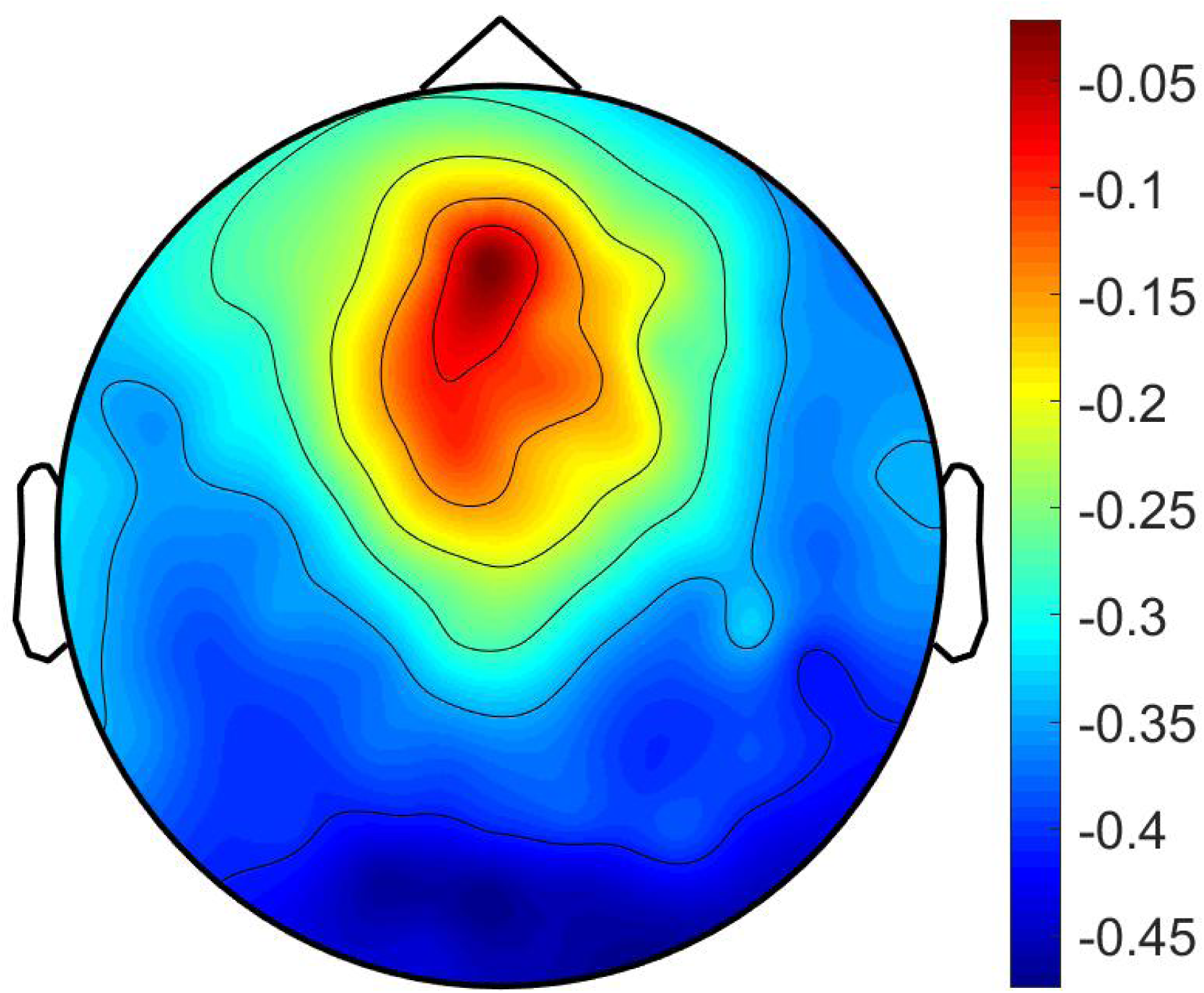
represents Sensor topography of correlation between peak alpha frequency and age. Colorbar represents Spearman’s rank correlation value.

**Supplementary Table 1.** Statistical table for power, coherence and metastability measures. F-values, beta coefficients and goodness of fit for linear regression based analysis are reported.

|  | *Measure* | F(1,648)(p=value) | beta1 | R^2 |
| --- | --- | --- | --- | --- |
| POWER | Delta Power | 2.61(0.1) | 0.008(0.1) | 0.004 |
|  | Theta Power | 0.546(0.46) | 0.004(0.46) | 0.0008 |
|  | Alpha Power | 0.08(0.76) | 0.0002(0.76) | 0.0001 |
|  | Beta Power | 41.3(<0.0001) | 0.03(<0.0001) | 0.06 |
| COHERENCE | Global Coherence(Delta) | 22(<0.0001) | 0.0005(<0.0001) | 0.03 |
|  | Global Coherence(Theta) | 6.76(<0.0001) | 0.0002(<0.009) | 0.01 |
|  | Global Coherence(Alpha) | 60(<0.0001) | -0.0008(<0.0001) | 0.084 |
|  | Global Coherence(Beta) | 0.03(0.86) | ~0(0.86) | ~0 |
| METASTABILITY | Metastability(Delta) | 95(<0.0001) | 0.0004(<0.0001) | 0.128 |
|  | Metastability(Theta) | 105(<0.0001) | 0.0004(<0.0001) | 0.139 |
|  | Metastability(Alpha) | 77.5(<0.0001) | 0.0003(<0.0001) | 0.107 |
|  | Metastability(Beta) | 21.6(<0.0001) | 0.0001(<0.0001) | 0.032 |
|  | Peak Alpha Frequency | 109(0.0001) | -0.01(<0.0001) | 0.144 |
|  | Alpha-Beta topographical segregation index | 65.7(<0.0001) | 0.003(<0.0001) | 0.09 |

**Supplementary Table 2.** Tabulates the between group test for beta power. 10000 iterations were performed to generate surrogate data for each comparison. Reported values correspond to p-values, effect size, group difference in means in that order.

| Beta Power | YA | ME | ML | OA |
| --- | --- | --- | --- | --- |
| YA |  | <0.001, -0.48, -0.9679 | <0.001, -0.61, -1.28 | <0.001, -0.83, -1.73 |
| ME |  |  | <0.21, -0.14, -0.3 | <0.0016, -0.34, -0.76 |
| ML |  |  |  | <0.06, -0.19, -0.4496 |

**Supplementary Table 3.** Peak Alpha Frequency(PAF) Categorical Analysis. Tabulates the between group test for peak alpha frequency(PAF). 10000 iterations were performed to generate surrogate data for each comparison. Reported values correspond to p-values, effect size, group difference in means in that order.

| PAF | YA | ME | ML | OA |
| --- | --- | --- | --- | --- |
| YA |  | 0.02, 0.2674, -0.193 | <0.001, -0.61, -1.28 | <0.001, -0.83, -1.73 |
| ME |  |  | 0.21, -0.14, -0.3 | 0.0016, -0.34, -0.76 |
| ML |  |  |  | 0.06, -0.19, -0.4496 |

\*p-value, effect size, group difference in means

**Supplementary Table 4.** Alpha-Beta Topographical segregation. Tabulates the between group test for alpha-beta segregation measure. 10000 iterations were performed to generate surrogate data for each comparison. Reported values correspond to p-values, effect size, group difference in means in that order.

Alpha-Beta Topographical segregation
| ANG_SEP | YA | ME | ML | OA |
| --- | --- | --- | --- | --- |
| YA |  | 0.03 , -0.25, -2.81 | <0.001, -0.47, -5.55 | <0.001, -0.77, -9.06 |
| ME |  |  | 0.03 , -0.24, -2.7 | <0.001, -0.55, -6.24 |
| ML |  |  |  | 0.0064, -0.289, -3.5 |

**Supplementary Table 5.** Global Coherence Categorical Analysis. Tabulates the between group test for global coherence measure. 10000 iterations were performed to generate surrogate data for each comparison. Reported values correspond to p-values, effect size, group difference in means in that order.

| Delta | YA | ME | ML | OA |  | Theta | YA | ME | ML | OA |
| --- | --- | --- | --- | --- | --- | --- | --- | --- | --- | --- |
| YA |  | 0.88 ,-0.01, 0.0006 | 0.24, -0.15 -0.006 | 0.0003, -0.48, -0.02 |  | YA |  | 0.6 ,-0.06, 0.001 | 0.43,0.09,0.002 | 0.05, -0.25, -0.009 |
| ME |  |  | 0.21 ,-0.14, -0.006 | <0.001, -0.43, -0.02 |  | ME |  |  | 0.8,0.02, 0.0009 | 0.01,-0.26,-0.011 |
| ML |  |  |  | 0.011,-0.28, -0.01 |  | ML |  |  |  | 0.01,-0.29,-0.01 |
| Alpha | YA | ME | ML | OA |  | Beta | YA | ME | ML | OA |
| YA |  | 0.1 ,0.18, 0.01 | <0.001, 0.63,0.03 | <0.001, 0.73, 0.03 |  | YA |  | 0.36,0.1,0.002 | 0.03, 0.25, 0.005 | 0.78,0.031, 0.0007 |
| ME |  |  | 0.0002 ,0.43, 0.02 | <0.001, 0.52, 0.02 |  | ME |  |  | 0.16,0.15,0.003 | 0.52,-0.06,-0.001 |
| ML |  |  |  | 0.4, 0.08, 0.003 |  | ML |  |  |  | 0.04,-0.2,-0.005 |

**Supplementary Table 6.** Metastability Categorical Analysis. Tabulates the between group test for metastability measure. 10000 iterations were performed to generate surrogate data for each comparison. Reported values correspond to p-values, effect size, group difference in means in that order.

| Delta | YA | ME | ML | OA |  | Theta | YA | ME | ML | OA |
| --- | --- | --- | --- | --- | --- | --- | --- | --- | --- | --- |
| YA |  | 0.007,-0.32, -0.004 | <0.001, -0.66, 0.01 | <0.001, -1.0, -0.01 |  | YA |  | <0.001 ,-0.41, -0.006 | <0.001,-0.74,-0.01 | <0.001, -1.15, -0.02 |
| ME |  |  | 0.002 ,-0.33, -0.006 | <0.001, -0.74, -0.01 |  | ME |  |  | 0.003 ,-0.33, -0.006 | <0.001, -0.73, -0.01 |
| ML |  |  |  | 0.0006, -0.39, -0.008 |  | ML |  |  |  | <0.001, -0.4, -0.009 |
| Alpha | YA | ME | ML | OA |  | Beta | YA | ME | ML | OA |
| YA |  | 0.02 ,-0.27, -0.003 | <0.001, -0.62, -0.009 | <0.001, -0.96, -0.01 |  | YA |  | 0.04 ,-0.24, -0.003 | 0.03, -0.26, -0.003 | <0.001, -0.6, -0.009 |
| ME |  |  | 0.004 ,0.32, 0.005 | <0.001, -0.67, -0.01 |  | ME |  |  | 0.82 ,-0.025, -0.0004 | 0.002, -0.33, -0.006 |
| ML |  |  |  | 0.0003, -0.38, -0.007 |  | ML |  |  |  | 0.006, -0.29, -0.005 |

**Fig S3.**
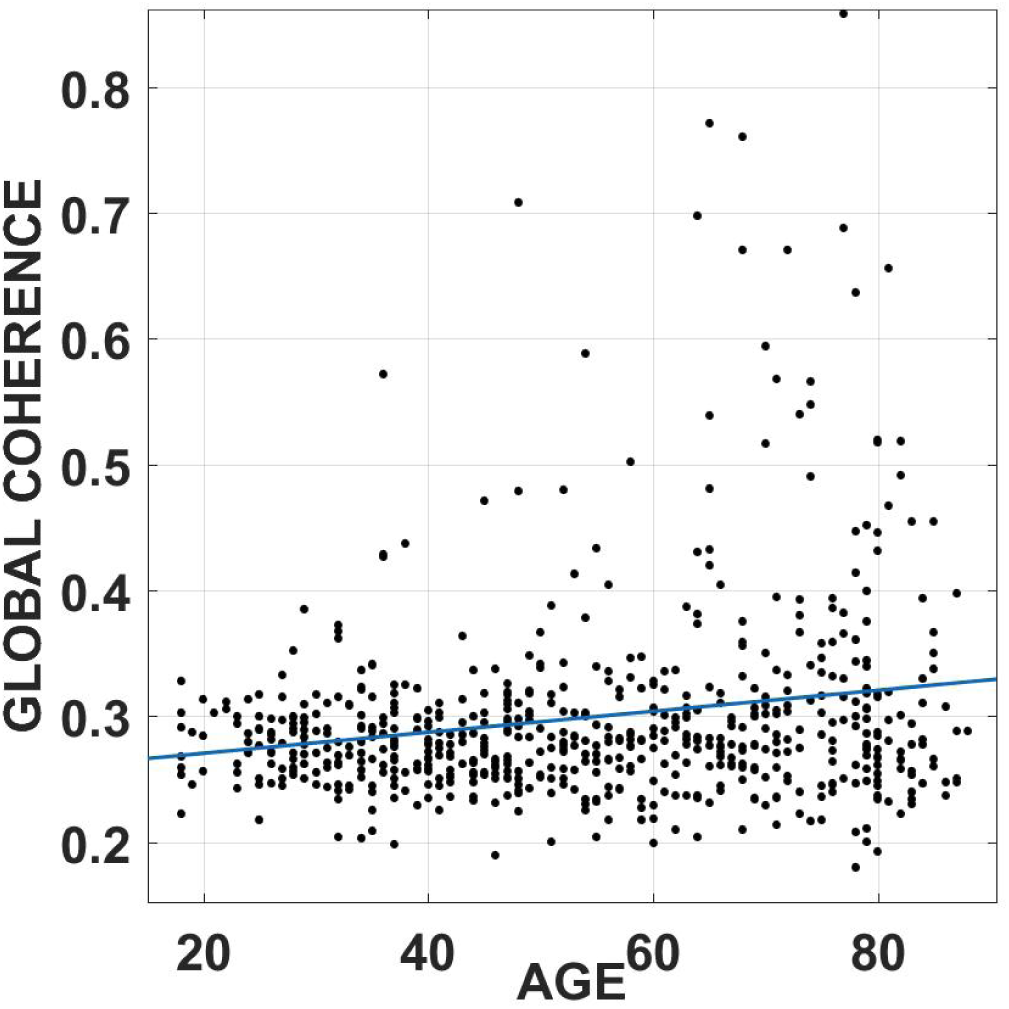
represents a scatter plot of Global Coherence in theta band(3-7Hz) (ICA CORRECTED) as a function of age. ECG and EOG signals were subtracted from the data using an automated ICA procedure as outlined in the main text.

**Fig. S4.**
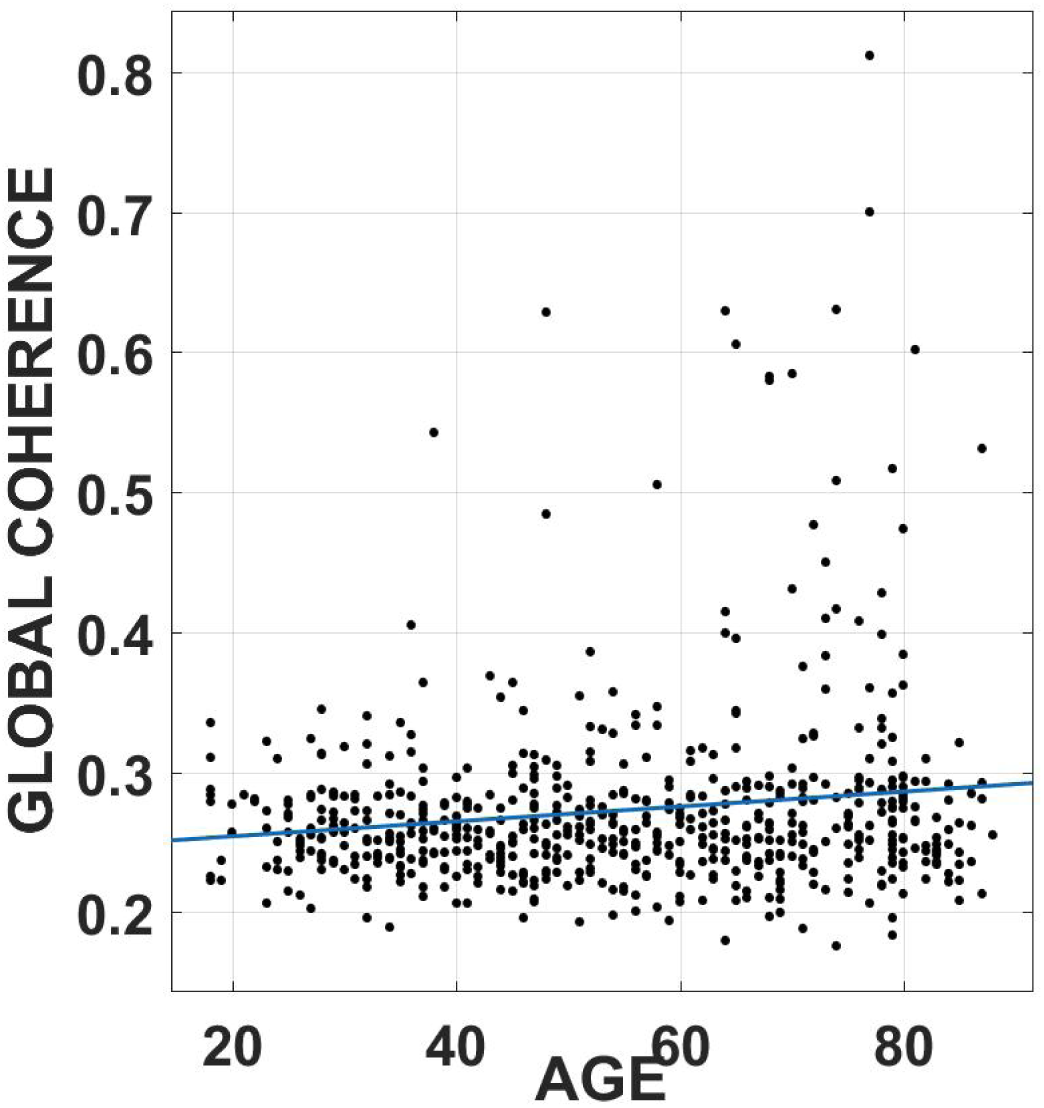
represents a scatter plot of Global Coherence in alpha band(8-12Hz) (ICA CORRECTED) as a function of age. ECG and EOG signals were subtracted from the data using an automated ICA procedure as outlined in the main text.

**Fig. S5.**
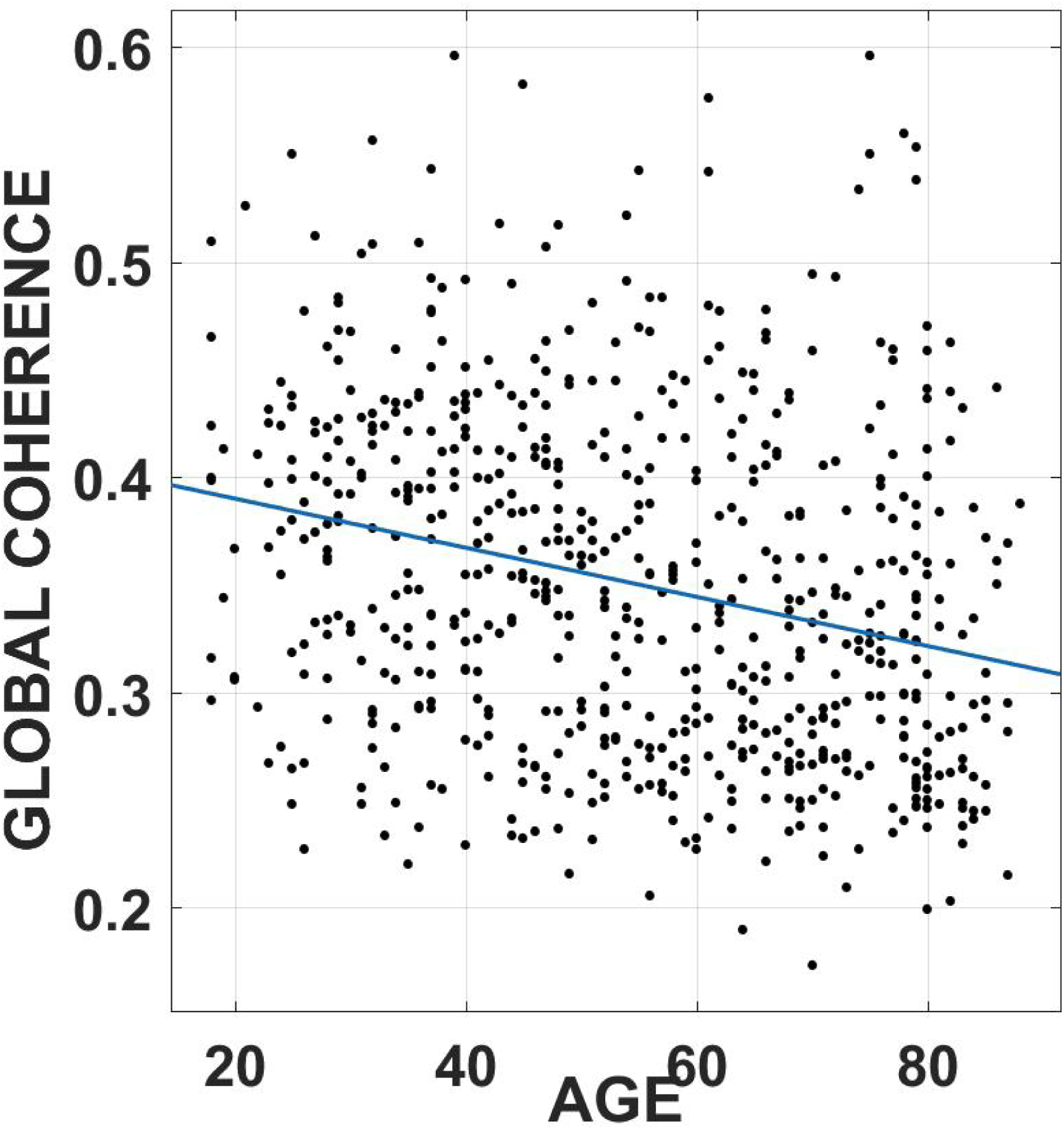
represents a scatter plot of Global Coherence in alpha band(8-12Hz) (ICA CORRECTED) as a function of age. ECG and EOG signals were subtracted from the data using an automated ICA procedure as outlined in the main text.

**Fig. S6.**
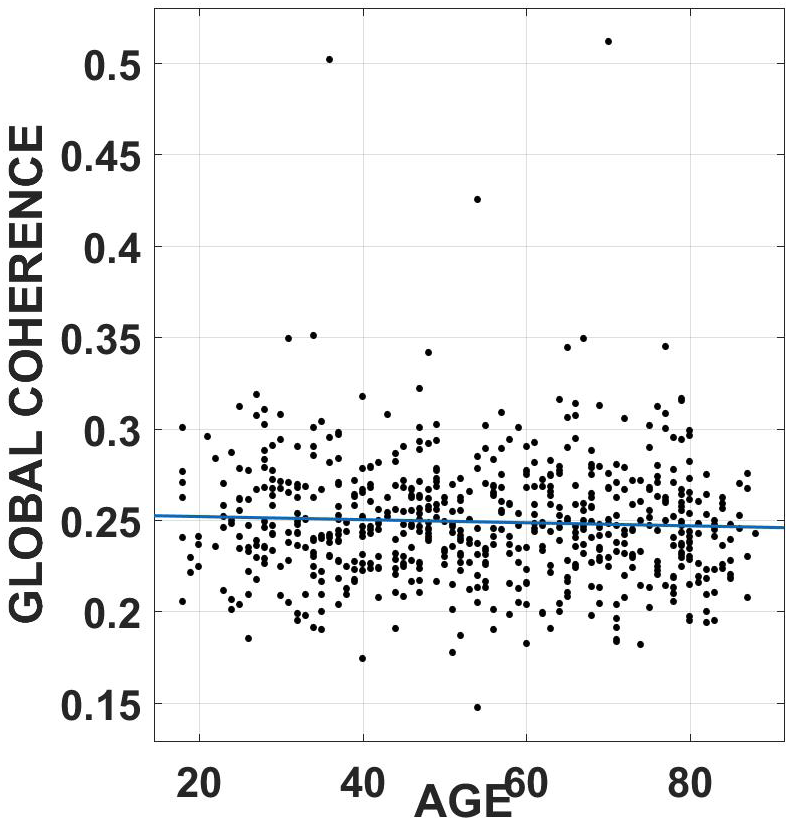
represents a scatter plot of Global Coherence in beta band(16-25Hz) (ICA CORRECTED) as a function of age. ECG and EOG signals were subtracted from the data using an automated ICA procedure as outlined in the main text.

**Fig. S7.**
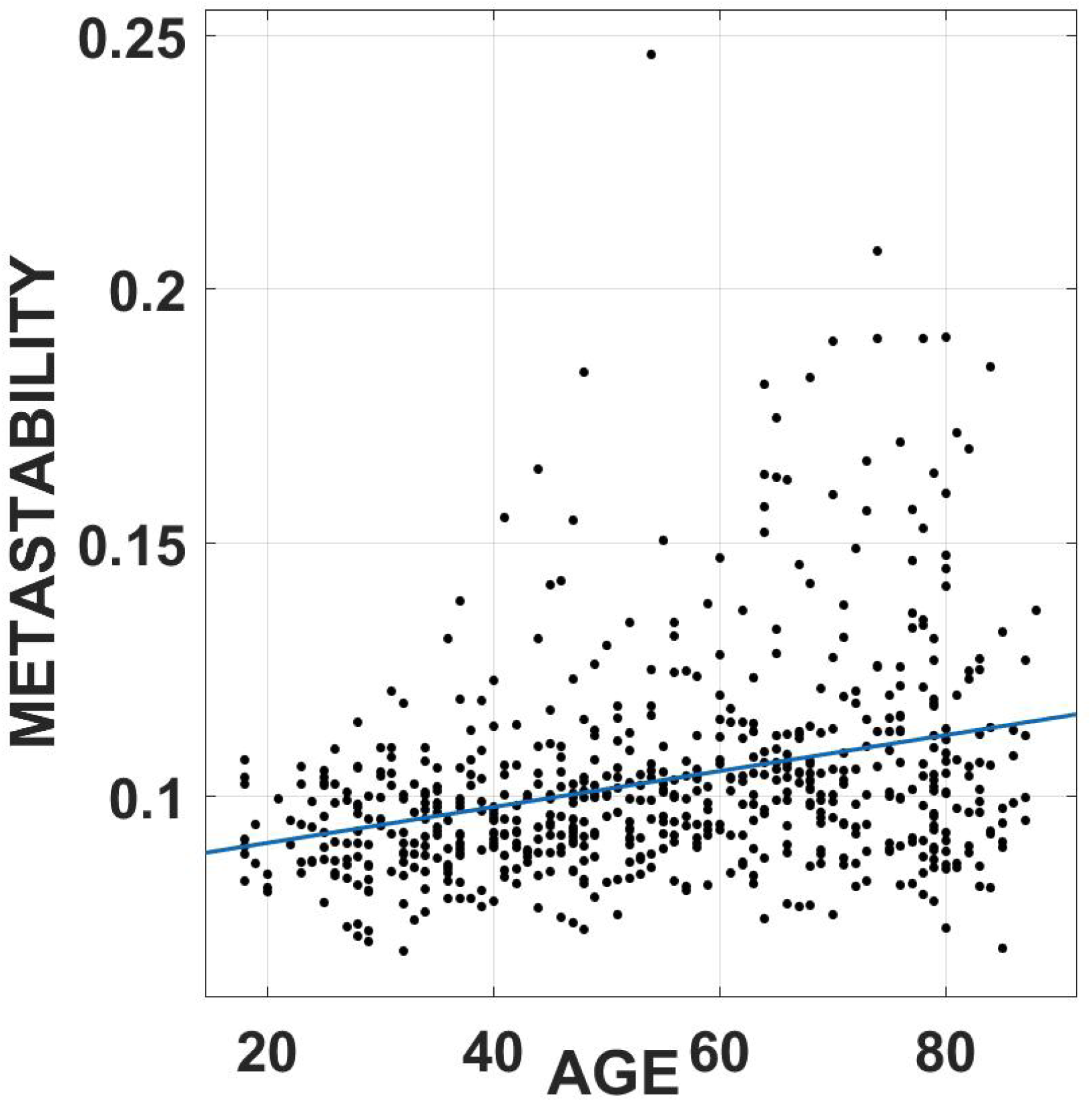
Represents a scatter plot of metastability in delta band(1-3Hz) (ICA CORRECTED) as a function of age. ECG and EOG signals were subtracted from the data using an automated ICA procedure as outlined in the main text.

**Fig. S8.**
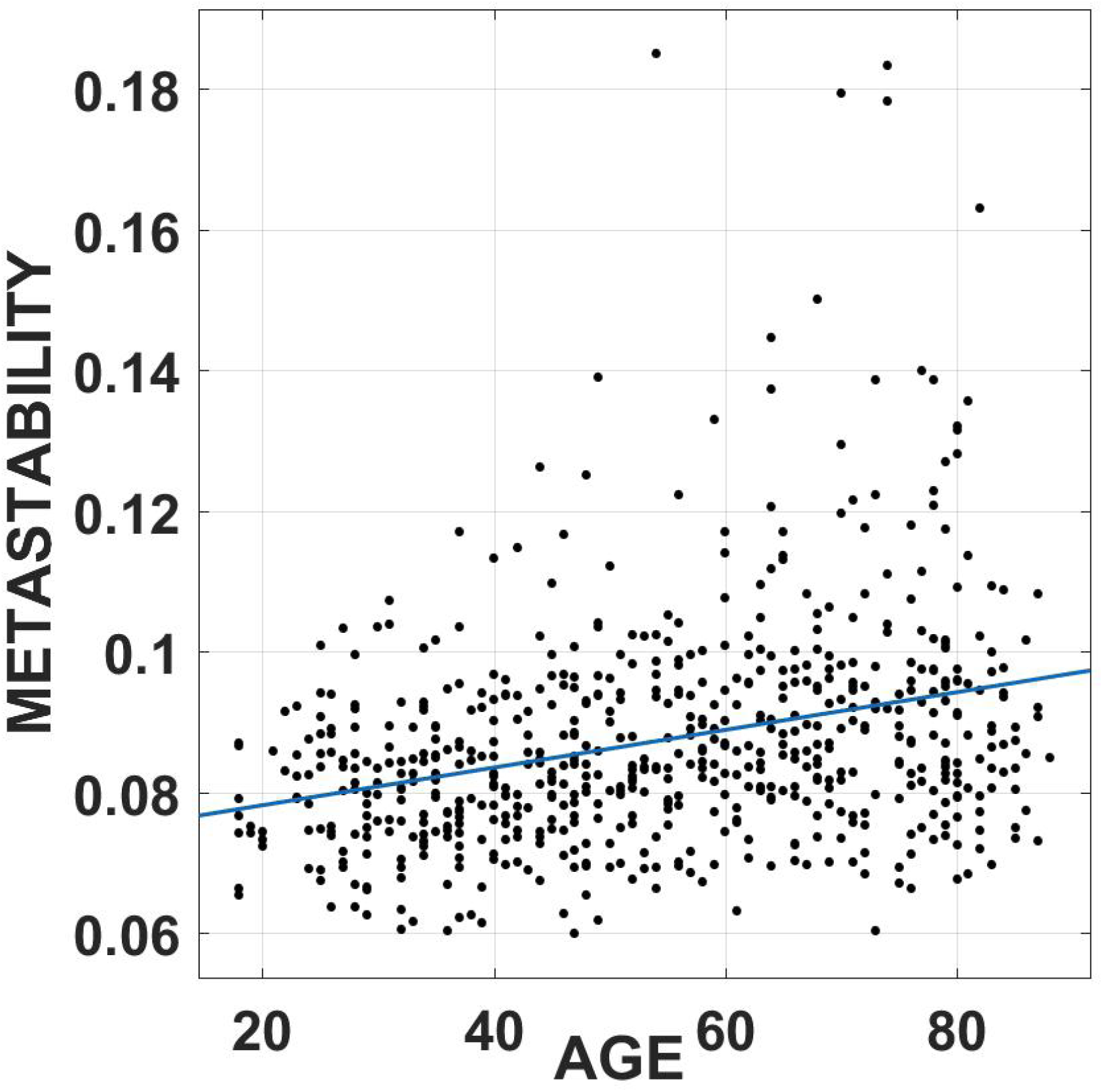
represents a scatter plot of metastability in alpha band(8-12Hz) (ICA CORRECTED) as a function of age. ECG and EOG signals were subtracted from the data using an automated ICA procedure as outlined in the main text.

**Fig. S9.**
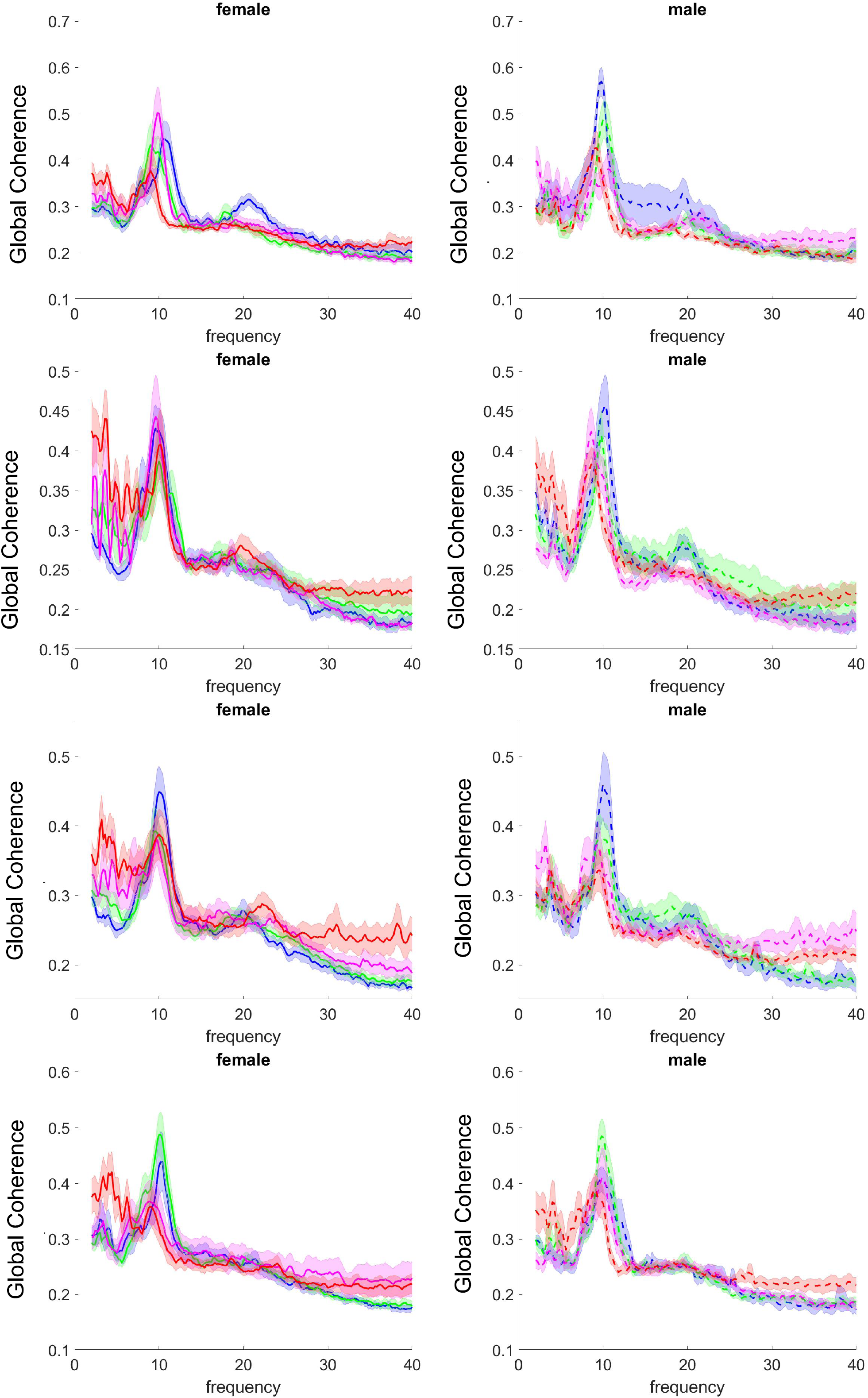
represents Gender-wise comparison of randomly sampled age groups. 50 samples were drawn at random from each age-group and GC calculated.

**Fig. S10.**
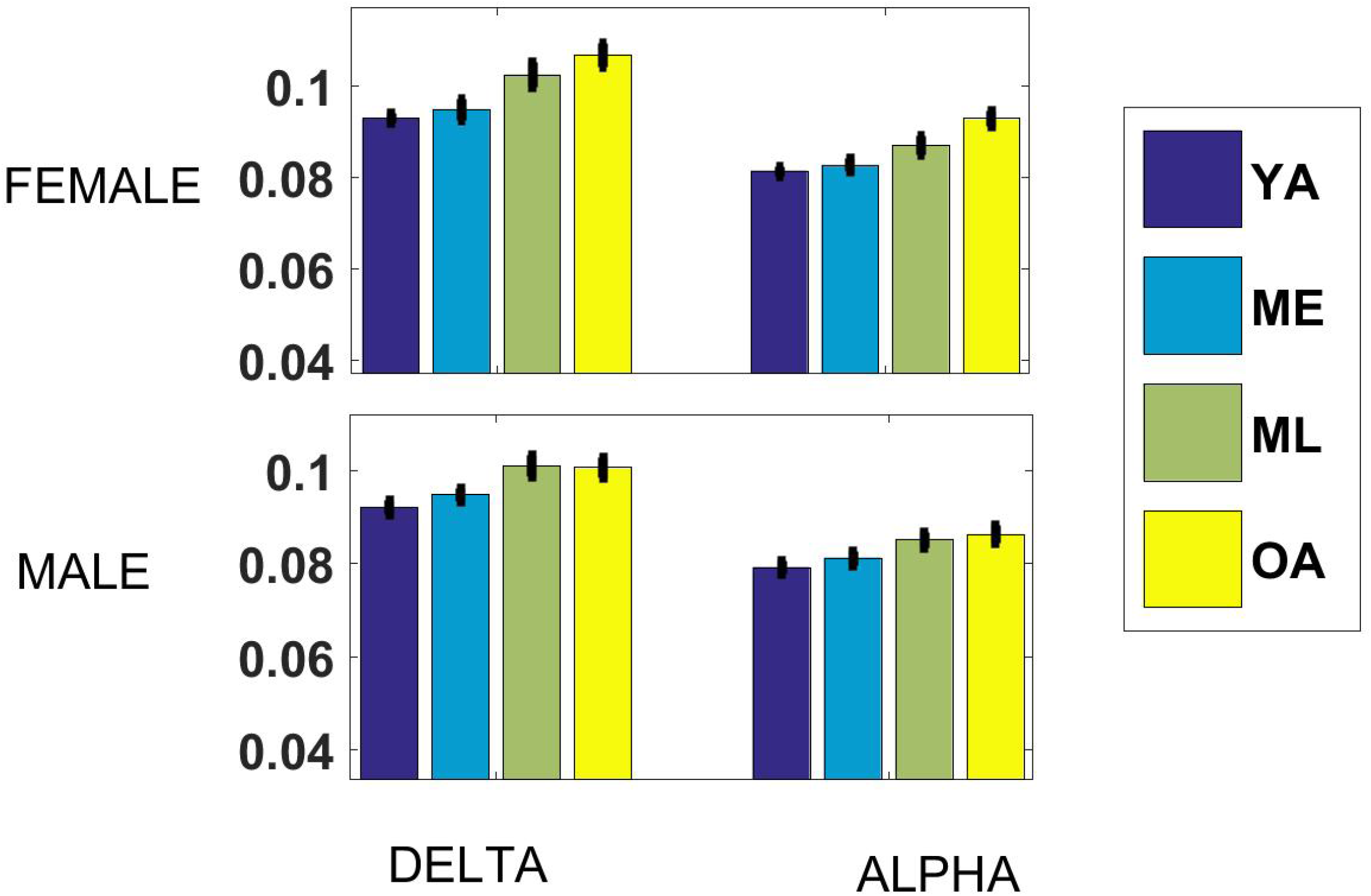
Gender-wise metastability in delta and alpha band. Data consisted of 328 Males and 322 Females. Bar plots represent metastability in the delta and theta bands for the 4 age groups.

**Supplementary Table 7.** Rho and P-val for partial correlation between precision in VSTM task and frequency-specific coherence and metastability(age regressed out).

| GLOBAL COHERENCE |  |  |
| --- | --- | --- |
| BAND | RHO | P |
| DELTA | -0.0018 | 0.96 |
| THETA | 0.01 | 0.72 |
| ALPHA | 0.09 | 0.01 |
| BETA | 0.08 | 0.06 |
| METASTABILITY |  |  |
| BAND | RHO | P |
| DELTA | 0.04 | 0.27 |
| THETA | 0.04 | 0.24 |
| ALPHA | 0.03 | 0.03 |
| BETA | 0.01 | 0.79 |

